# Spliceosomic dysregulation unveils *NOVA1* as an actionable therapeutic target in pancreatic neuroendocrine tumors

**DOI:** 10.1101/2022.02.09.479525

**Authors:** Sergio Pedraza-Arevalo, Emilia Alors-Pérez, Ricardo Blázquez-Encinas, Aura D. Herrera-Martínez, Juan M. Jiménez-Vacas, Antonio C. Fuentes-Fayos, Óscar Reyes, Sebastián Ventura, Rafael Sánchez-Sánchez, Rosa Ortega-Salas, Raquel Serrano-Blanch, María A. Gálvez-Moreno, Manuel D. Gahete, Alejandro Ibáñez-Costa, Raúl M. Luque, Justo P. Castaño

## Abstract

Dysregulation of the splicing machinery is emerging as a hallmark in cancer due to its association with multiple dysfunctions in tumor cells. Inappropriate function of this machinery can generate tumor-driving splicing variants and trigger oncogenic actions. However, its role in pancreatic neuroendocrine tumors (PanNETs) is poorly defined. In this study we aimed to characterize the expression pattern of a set of splicing machinery components in PanNETs, and their relationship with aggressiveness features. A qPCR-based array was first deployed to determine the expression levels of components of the major (n=13) and minor spliceosome (n=4) and associated splicing factors (n=27), using a microfluidic technology in 20 PanNETs and non-tumoral adjacent samples. Subsequently, *in vivo* and *in vitro* models were applied to explore the pathophysiological role of *NOVA1*. Expression analysis revealed that a substantial proportion of splicing machinery components was altered in tumors. Notably, key splicing factors were overexpressed in PanNETs samples, wherein their levels correlated with clinical and malignancy features. Using *in vivo* and *in vitro* assays, we demonstrate that one of those altered factors, *NOVA1*, is tightly related to cell proliferation, alters pivotal signaling pathways and interferes with responsiveness to drug treatment in PanNETs, suggesting a role for this factor in the aggressiveness of these tumors and its suitability as therapeutic target. Altogether, our results unveil a severe alteration of the splicing machinery in PanNETs and identify the putative relevance of *NOVA1* in tumor development/progression, which could provide novel avenues to develop diagnostic biomarkers and therapeutic tools for this pathology.

## Introduction

Neuroendocrine tumors (NETs) comprise a highly heterogeneous group of neoplasms that arise from cells of the diffuse neuroendocrine system and are thus widely distributed throughout the body, being more common in gastroenteropancreatic (GEP) and bronchopulmonary tracts (1). Pancreatic NETs (PanNETs) mainly derive from the hormone-producing cells of the pancreas and represent less than 3 % of pancreatic primary neoplasms but show a rapid increase in recent years (1-3). PanNETs diagnosis is hampered by a combined lack of specific symptoms and useful biomarkers, which often leads to late detection of tumors already displaying high grade and metastasis (1, 2). In addition, PanNET heterogeneity, derived from tumor-specific molecular features, hinders their clinical management and pharmacological treatment (4), which primarily employ somatostatin analogs and inhibitors of mTOR pathway and angiogenesis. These would benefit from precision medicine approaches, yet specific response-predictive biomarkers and broadly actionable targets are still lacking (4, 5).

Recent studies have analyzed in detail the molecular features of PanNETs, defining their genomic, epigenomic and transcriptomic landscapes and unveiling key mutations and alterations that contribute to tumorigenesis (2, 6-8). The most frequent mutations in PanNETs involve *MEN1* and other genes related to chromatin remodeling, core genes of the PI3K/mTOR pathway (e.g., *TSC1, TSC2* and *PTEN*), and DNA repair (*MUTYH, BRCA2*) and cell cycle machineries, as well as mutually exclusive mutations in *ATRX* and *DAXX*, which are related with alternative lengthening of telomeres (ALT) (8, 9). Likewise, epigenetic alterations including DNA methylation and miRNA dysregulations have been explored and shown to frequently converge upon elements comprising the same regulatory pathways (2, 7). On the other hand, the potential role of an emergently relevant regulatory mechanism in cancer, i.e. alternative splicing (10), in PanNETs is poorly known.

Earlier evidence uncovered the overexpression in GEPNETs of aberrantly spliced, truncated variants of somatostatin receptor subtype 5, SST_5_TMD4 (11), and ghrelin gene, named In1-ghrelin (12), which have been associated with oncogenesis and aggressiveness in various hormone-related cancers (13-15). These observations were in line with the growing notion that altered splicing represents a genuine transversal hallmark influencing all classic cancer hallmarks (10, 16) and prompted us to explore the underlying mechanisms. Aberrant splicing may result from an incorrect intron-exon processing of the pre-RNAs into mature RNAs by the spliceosome, a macromolecular machinery composed by five small nuclear ribonucleoproteins (snRNPs; formed by small nuclear RNAs, snRNAs, and associated proteins) working in concert with more than 150 proteins termed splicing factors (10, 16-18). Recently, dysregulation of the splicing machinery, by mutations or altered expression of their specific elements, is increasingly regarded as a pathological trigger for tumor development and treatment resistance (10, 16-18). Indeed, our group has identified specific factors altered in various cancers, from pancreatic adenocarcinoma to glioblastoma, prostate cancer, hepatocarcinoma, or pituitary neuroendocrine tumors, where they play distinct oncogenic roles (19-23). However, to the best of our knowledge, the potential alteration of the splicing machinery in PanNETs and its possible relation with tumor biology has not been reported to date.

Thus, in this study we aimed to define the expression profile of a selected set of key components of the splicing machinery in PanNETs and evaluate their possible alterations and associations with clinical parameters. Furthermore, we plan to elucidate the functional role of particularly dysregulated factors using *in vitro* and *in vivo* PanNETs models, with the ultimate goal of identifying potential biomarkers and/or actionable targets to improve patient diagnosis and treatment.

## Material and methods

### Patients and samples

Formalin fixed paraffin-embedded samples (FFPE, n = 20) were obtained from primary PanNETs (NCNN-guidelines) (24), summarized in **Table 1**. Non-tumoral adjacent tissue, used as reference-control, was extracted from the same piece and both tissues were separated by expert pathologists. Patients were managed following current recommendations and guidelines. Patients were classified according to the ENETS and WHO criteria (tumor site and size, Ki67 index, necrosis, mitotic rate and relapse of the disease) (25). This study was conducted in accordance with the Declaration of Helsinki and approved by the Reina Sofia University Hospital (Córdoba, Spain) Ethics Committee. Written informed consents from patients were obtained through the Andalusian Biobank (Servicio Andaluz de Salud, application code S1900499).

**Table 1.** Summary of clinical parameters of the PanNETs patients. Overall clinical and demographic data of patients that participated in this study.

|  |  |
| --- | --- |
| Number of samples | 20 |
| Age (mean) | 55 ± 14 years |
| Body Mass Index | 28.00 ± 3.48 kg/m<sup>2</sup> |
| Gender (female) | 57.1 % |
| Gender (male) | 42.9 % |
| Smoking | 68.8 % |
| Family history of neoplasia | 12.5 % |
| Comorbidities | 83.3 % |

### Cell culture

Functional assays were carried out in two PanNETs cell lines. Specifically, we used carcinoid derived BON-1 cells (26) and somatostatinoma derived QGP-1 cells (27). BON-1 cells were cultured in Dulbecco’s Modified Eagles Medium complemented with F12 (DMEM-F12; Life Technologies, Barcelona, Spain) and QGP-1 cells were grown in RPMI 1640 (Lonza, Basel, Switzerland), both supplemented with 10 % fetal bovine serum (FBS; Sigma-Aldrich, Madrid, Spain), 1 % glutamine (Sigma-Aldrich) and 0.2 % antibiotic (Gentamicin/Amphotericin B; Life Technologies). Both cell lines were grown at 37 °C, in a humidified atmosphere with 5.0 % CO_2_ and were verified for mycoplasma contamination as previously reported (28).

### Immunohistochemical analysis

Immunohistochemistry (IHC) technique was performed to study the protein expression levels of NOVA1 in FFPE PanNETs samples, using standard procedures (29) with a commercial antibody (#PA5-18895, Thermo Fisher, Waltham, MA, USA), checking the absence of staining in a negative control and a positive staining in a high expression tissue. An expert pathologist carried out a histopathologic analysis of the sections, following a blinded protocol, indicating +, ++ and +++ as low, moderate and high staining intensity, respectively of tumoral compared to non-tumoral adjacent tissue.

### Modulation of *NOVA1* expression by specific siRNA and overexpression plasmid

QGP-1 and BON-1 cells were transfected with a specific siRNA or overexpression plasmid, previously validated in our laboratory (SC111659 and SR303213, respectively; Origene, Rockville, MD, USA). Specifically, cells were seeded in 6-well culture plates and transfected with *NOVA1* siRNA (100 nM) or plasmid (1 µg), using Lipofectamine RNAiMAX or Lipofectamine 2000 Transfection Reagents (Invitrogen, Thermo Fisher), respectively, following manufacturer instructions. Scramble siRNA and empty vector (mock) were used as control. Success of the silencing was validated by qPCR and western blot.

### Alamar Blue proliferation assay

Alamar Blue fluorescent assay (Life Technologies) was used to determine cell proliferation as previously reported (30). In all instances, cells were seeded per quadruplicate and all assays were repeated a minimum of three times.

### RNA isolation and retrotranscription

Total RNA from cell lines was isolated using TRIzol Reagent (Sigma-Aldrich) treated with DNase (Promega, Barcelona, Spain) following manufacturer instructions. Regarding FFPE samples, RNA was isolated RNeasy FFPE Kit (Qiagen, Limburg, Netherlands) according to the manufacturer protocol. The amount of RNA recovered was determined using the NanoDrop2000 spectrophotometer (Thermo Fisher) and its quality was assessed by the same system using the Absorbance Ratio A260/280 and A260/230, requiring a minimum of 1.8 in both. One µg of RNA was reverse transcribed to cDNA using random hexamer primers with the First Strand Synthesis Kit (Thermo Fisher).

### Quantitative real time PCR (qPCR)

qPCR reactions were carried out using the Stratagene Mx3000p system with the Brilliant III SYBR Green Master Mix (Stratagene, La Jolla, CA, USA) as previously described (21, 23). Specifically, *MKI67, CCND1, CASP3, TP53, Δ133TP53, TERT, TERT tv1, ATRX* and *DAXX* expression levels were analyzed. Primers are listed in **Supplemental Table 1** (*Δ133TP53* (31), *TERT*, and *TERT tv1* (32)). Results were validated as previously reported (33), normalizing all genes with a normalization factor, calculated from expression values of *ACTB, GAPDH* and *HPRT1* housekeeping genes, using Genorm Software.

### qPCR Dynamic Array

A Dynamic Array (Fluidigm, South San Francisco, CA, USA), based on microfluidic technique for gene expression analysis, as previously reported in detail (21, 23).

### Western Blot

QGP-1 and BON-1 transfected with siRNAs were lysed to analyze protein expression and phosphorylation by western blot, using standard procedures (13) and specific antibodies for ERK (sc-154, Santa Cruz Biotechnology, Dallas, TX, USA); phospho-ERK (#4370S, Cell Signaling, Beverly, MA, USA); AKT (#9272S, Cell Signaling); phosphor-AKT (#4060S); phospho-PDK1 (#3061, Cell Signaling); phospho-PTEN (#9551, Cell Signaling); phospho-p53 (#9284S, Cell Signaling); ATRX (HPA001906, Sigma Aldrich); DAXX (HPA008736, Sigma Aldrich); and NOVA1 (#PA5-18895, Thermo Fisher); as well as with the appropriate secondary antibodies: HRP-conjugated goat-anti rabbit (#7074s; Cell Signaling) or rabbit anti-goat (#31753; Thermo Fisher) IgG. A densitometry analysis of the bands obtained was carried out with ImageJ software (version 1.5, developed by NIH, Bethesda, MD, USA), using total protein or Ponceau as normalizing factor of correspondent phosphorylated protein.

### Xenograft model

Animal maintenance and experiments were carried out following the European Regulation for Animal Care and under the approval of the University of Córdoba Research Ethics Committee. Seven-week-old male athymic BALB/cAnNRj-Foxn1nu mice (Janvier Labs, Le Genest-Saint-Isle, France; n = 4 mice), were subcutaneously grafted in the flank with 3 × 10^6^ BON-1 cells transfected with mock and *NOVA1* plasmids in each flank, resuspended in 100 µl of basement membrane extract (29). Tumor growth was monitored twice per week for 5 weeks, by using a digital caliper.

### Statistical analyses

Data were first evaluated for parametric distribution with Kolmogorov–Smirnov test and were expressed as mean ± SEM (standard error of the mean) or median plus interquartile range for human data. Statistical comparisons between groups were performed by unpaired parametric t-test and non-parametric Mann-Whitney U test, according to normality. Multiple comparisons of more than two groups were performed for analysis of variance (one-way ANOVA) followed by Dunnett’s test. Pearson’s or Spearman’s bivariate correlations were performed for quantitative variables, in case they were parametric or not, respectively. The receiver operating characteristic (ROC) curves were used to evaluate the suitability of genes to distinguish different groups of samples. Additional analyses enabled to check the ability of several factors to distinguish between tumoral and non-tumoral samples. Random forest and simple logistic regression analyses were carried out with R language and followed by cross-validation to select a group of factors with a good ability to make clusters with samples. The VIP score was performed through Metaboanalyst software 4.0 (34). Statistical analyses were assessed using GraphPad Prism 7 (GraphPad Software, La Jolla, CA, USA) and correlations were carried out using SPSS 22 (IBM SPSS Statistics Inc., Chicago, IL, USA). The p-values were two-sided and statistical significance was considered when p < 0.05.

## Results

### Splicing machinery is deeply dysregulated in PanNETs and associated with clinical parameters

To assess the expression profile of a selected set of components of the splicing machinery, we employed a qPCR Array based on microfluidics, and measured their mRNA levels in a cohort of 20 primary tumors from patients with PanNETs, comparing the tumor tissue with the non-tumoral adjacent tissue, used as reference. The set included major spliceosome (n = 13) and minor spliceosome (n = 4) elements, as well as a group of associated splicing factors (n = 27) that were selected based on the literature (**Supplemental Figure 1**). This approach revealed, for the first time, that seven components of major spliceosome, two of the minor spliceosome and ten splicing factors were upregulated, whereas only one splicing factor, *ESRP2*, was downregulated, in tumoral samples compared to control tissue (**Fig. 1A, Supplemental Figure 2**). These alterations accounted for nearly half of the components measured and included small nuclear RNAs (snRNAs), which comprise the core of the spliceosome. Interestingly, the potential relationship between the factors whose expression was found here to be altered is supported by the information already available in the literature, as it was shown by a STRING analysis (**Fig. 1B**).

**Figure 1.**
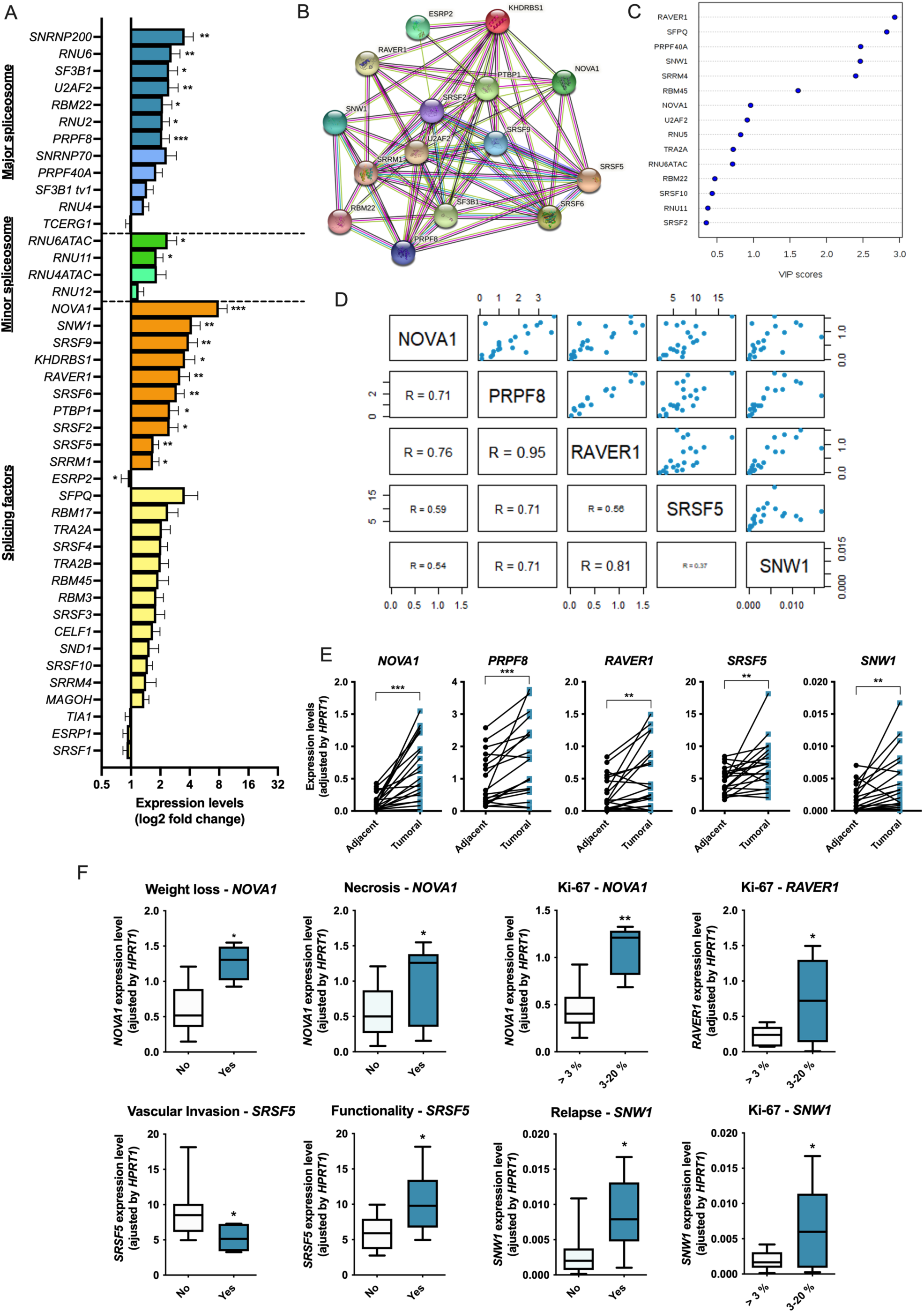
The components of the splicing machinery are clearly dysregulated in PanNETs and associated to clinical parameters. **A**. Expression levels of key components of the splicing machinery in PanNETs tumoral samples as compared to their paired non-tumoral adjacent tissue used as reference/control. The RNA levels were determined by qPCR and adjusted by *HPRT1* housekeeping gene. **B**. STRING analysis of relationships among altered components based on the information available in the literature. **C**. VIP score analysis showing the most modified factors in our cohort. **D**. Correlations between *NOVA1, PRPF8, RAVER1, SRSF5* and *SNW1* splicing machinery components. The upper-right panels represent the scatter plot of the correlations, and the lower-left panel represents the R of each correlation, the higher the size the better the correlation obtained. **E**. Paired comparison of these five components between tumoral and adjacent tissue. **F**. Associations between clinically relevant parameters of patient and tumors and the expression levels of the altered splicing machinery components selected on the basis of their higher changes in tumoral samples. Asterisks (*, p < 0.05; **, p < 0.01; ***, p < 0.001) indicate values that significantly differ from control. In all cases, data represent mean ± SEM or median plus interquartile range of n = 20 independent samples.

To classify these factors as possible biomarkers, we performed a principal component analysis (**Fig. 1C**), as well as a random forest and simple logistic regression analyses, which indicated the molecules that combinedly presented with the highest changes and best clustering features (**Supplemental Table 2**). Specifically, this approach identified five genes that stand out among all those measured: *NOVA1, PRPF8, RAVER1, SRSF5* and *SNW1*. Of note, these five splicing machinery components were found to correlate significantly and positively with each other in our cohort (**Fig. 1D**). A more detailed analysis revealed that they are, overall, highly overexpressed in tumoral tissue with respect to the non-tumoral adjacent reference tissue, and, interestingly, that the increased gene expression level was consistently observed in the vast majority of samples analyzed (**Fig. 1E**).

Additional analyses revealed that four out of the five selected factors displayed significant correlations with key parameters related to tumor functionality and patient prognosis, which may provide additional information on the importance of these alterations (**Fig. 1F**). Specifically, high levels of *NOVA1* gene expression showed a strong association with several parameters, including weight loss, necrosis of the primary tumor and Ki-67 index, this latter being typically linked to the proliferative status of the tumoral cells. Interestingly, this Ki-67 index was also associated with expression levels *RAVER1* and *SNW1*, suggesting a notable relationship between cell proliferation and altered splicing. Moreover, a high expression of *SNW1* was related also to relapse of the disease. Finally, a higher expression of *SRSF5* was linked to a higher presence of functionality but lower vascular invasion.

### The splicing factor *NOVA1* as a putative biomarker for PanNETs

The next step in our study was to assess the potential value of the five selected splicing machinery components as biomarkers for PanNETs. To this end, we made ROC curves for each component (**Fig. 2A**). These analyses showed that *NOVA1* expression exhibited the highest area under the curve (AUC), above 0.86, whereas the other four factors yielded AUC ranging between 0.65-0.75. These results prompted us to study this factor in more detail.

**Figure 2.**
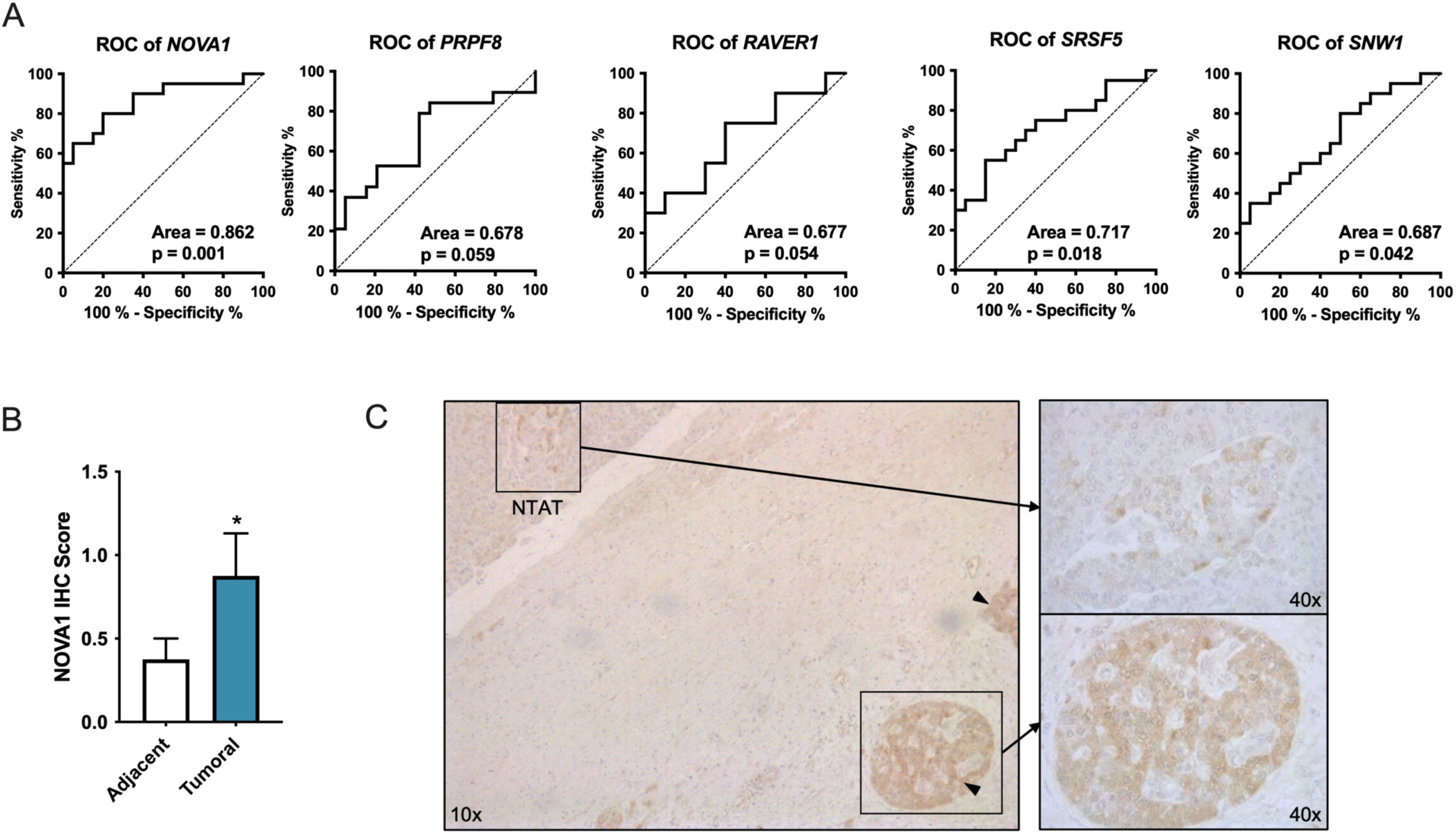
*NOVA1* as biomarker for PanNETs. **A**. Receiver operating characteristic (ROC) curve analysis was developed to determine the accuracy of *NOVA1, PRPF8, RAVER1, SRSF5* and *SNW1* expression to discriminate between tumoral and non-tumoral samples. **B**. Immunohistochemistry of NOVA1 protein was carried out in tissue section from PanNETs and staining in tumoral versus non-tumoral adjacent tissue was evaluated and scored by expert pathologists. **C**. Representative picture of a NOVA1 immunohistochemistry showing tumor and non-tumoral adjacent tissue (NTAT). Arrow heads point tumoral glands. Asterisks (*, p < 0.05) indicate values that significantly differ from control or between groups, respectively. In all cases, data represent mean ± SEM of n ≥ 3 independent samples.

Accordingly, we first applied an immunohistochemical analysis by expert pathologists, who confirmed the higher levels of NOVA1 protein in tumor tissue, compared with non-tumoral adjacent tissue (**Fig. 2B**). The staining of NOVA1 protein is clearly more prominent in the tumoral gland than in the non-tumoral adjacent tissue (NTAT), particularly in the endocrine tissue of the normal pancreas (Langerhans islets), which provides the most appropriate control tissue for PanNETs. Actually, this is one of the main limitations in the use of reference tissue in the study of PanNETs, and the results from immunohistochemistry help to overcome it.

### Overexpression of *NOVA1* increases cell proliferation and tumor growth

We next aimed to explore the possible functional role and mechanisms of action of this factor in PanNETs cells. To this end, we first tested the expression of *NOVA1* in two PanNETs model cell lines, QGP-1 and BON-1. We observed that both cell lines exhibited appreciable mRNA levels of *NOVA1*, which were high enough to perform silencing assays to examine the effect of *NOVA1* loss, but sufficiently moderate to also allow overexpression studies of this factor (**Fig. 3A**). Since we had observed that *NOVA1* was overexpressed in PanNETs, we initially overexpressed it in the two cell lines, and assessed functional features that could inform about tumor cell aggressiveness. Interestingly, in line with the previous results, *NOVA1* overexpression (**Supplemental Figure 3A**) increased cell proliferation in both cell lines, as measured by Alamar Blue assay, at different time points (24, 48, 72 h) after transfection (**Fig. 3B**).

**Figure 3:**
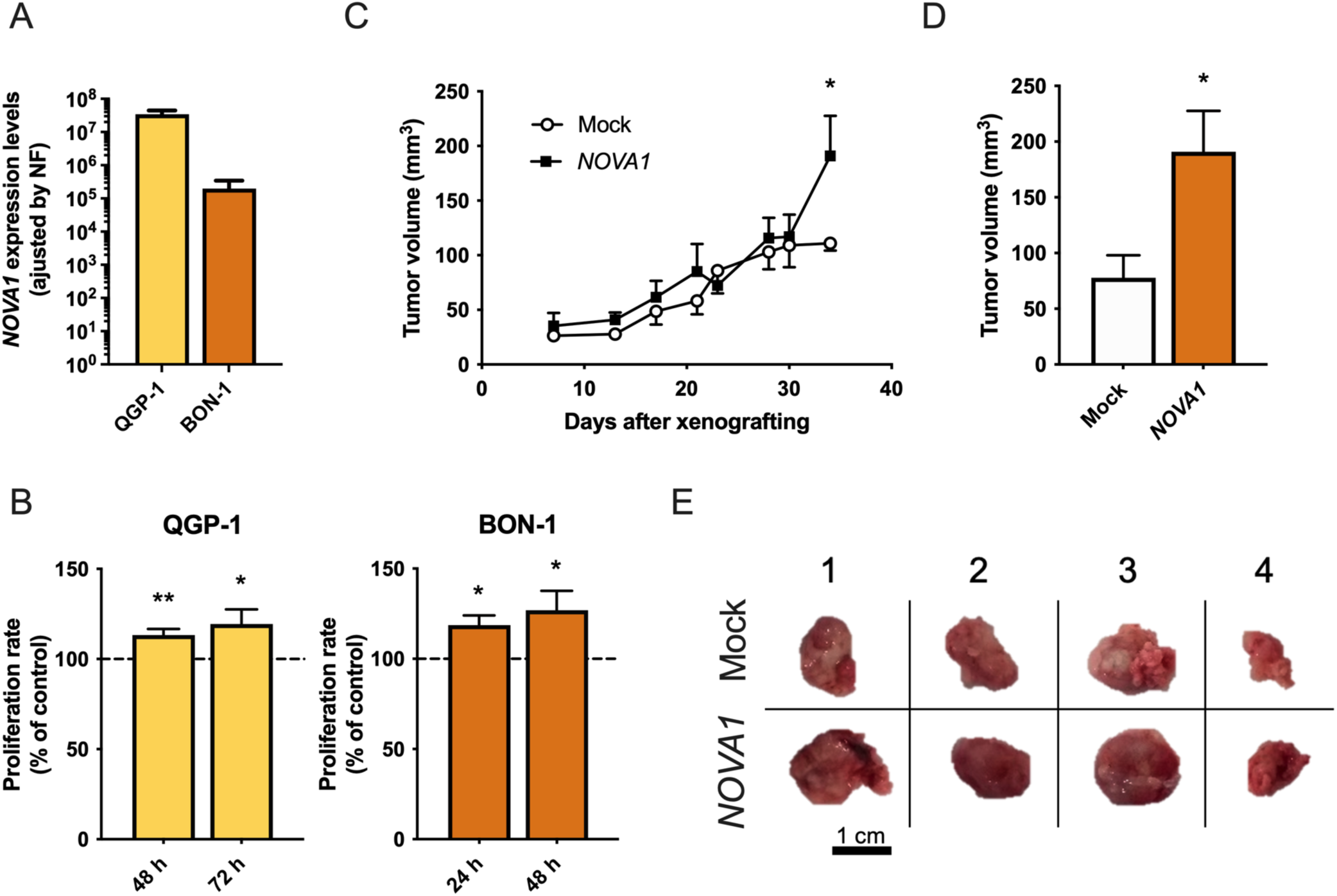
*NOVA1* is directly related with PanNETs cell proliferation. **A**. *NOVA1* expression levels in QGP-1 and BON-1 PanNETs cell lines as measured by qPCR. **B**. Proliferation rate of QGP-1 (yellow; left) and BON-1 (orange; right) at 48 and 72 h or 24 and 48 h, respectively, after *NOVA1* overexpression compared with mock plasmid transfection, used as control (100 %) and marked as tick line. **C**. BON-1 xenografted tumor growth in nude mice with *NOVA1* overexpression compared to mock. **D**. Comparison of tumor size at time of euthanasia; tumor volume is expressed as mm^3^. **E**. Picture of paired xenografted tumors with mock and *NOVA1* overexpression. Absolute mRNA levels were determined by qPCR and adjusted by normalization factor with *ACTB, GAPDH* and *HPRT1* housekeeping genes. Asterisks (*, p < 0.05; **, p < 0.01) indicate values that significantly differ from control. In all cases, data represent mean ± SEM of n ≥ 3 independent experiments.

Additionally, we developed a relevant preclinical model of PanNET xenograft tumors in mice, using BON-1 rather than QGP-1 cells, because the first showed a more aggressive phenotype. Thus, BON-1 cells transfected with *NOVA1*- or mock-plasmid were xenografted in nude mice. As shown in **Fig 3C-E**, BON-1 cells overexpressing *NOVA1* exhibited a higher growth rate over time than mock-control cells, producing larger tumors at the end of the experimental period. Taken together, these and the previous results strongly support the idea that the splicing factor *NOVA1* is directly related with cell proliferation in PanNETs and that increased levels of this factor may contribute to tumor aggressiveness.

### *NOVA1* as putative therapeutic target in PanNETs

Our next aim was focused on the study of *NOVA1* potential as a therapeutic target in these tumors. Given that this gene was overexpressed in the PanNETs samples, our experimental approach was to silence *NOVA1* expression in QGP-1 and BON-1 cell lines using a specific siRNA. Remarkably, this revealed that *NOVA1* silencing consistently decreased proliferation rate in both cell lines, compared to scramble siRNA, used as control (**Fig. 4A**).

**Figure 4.**
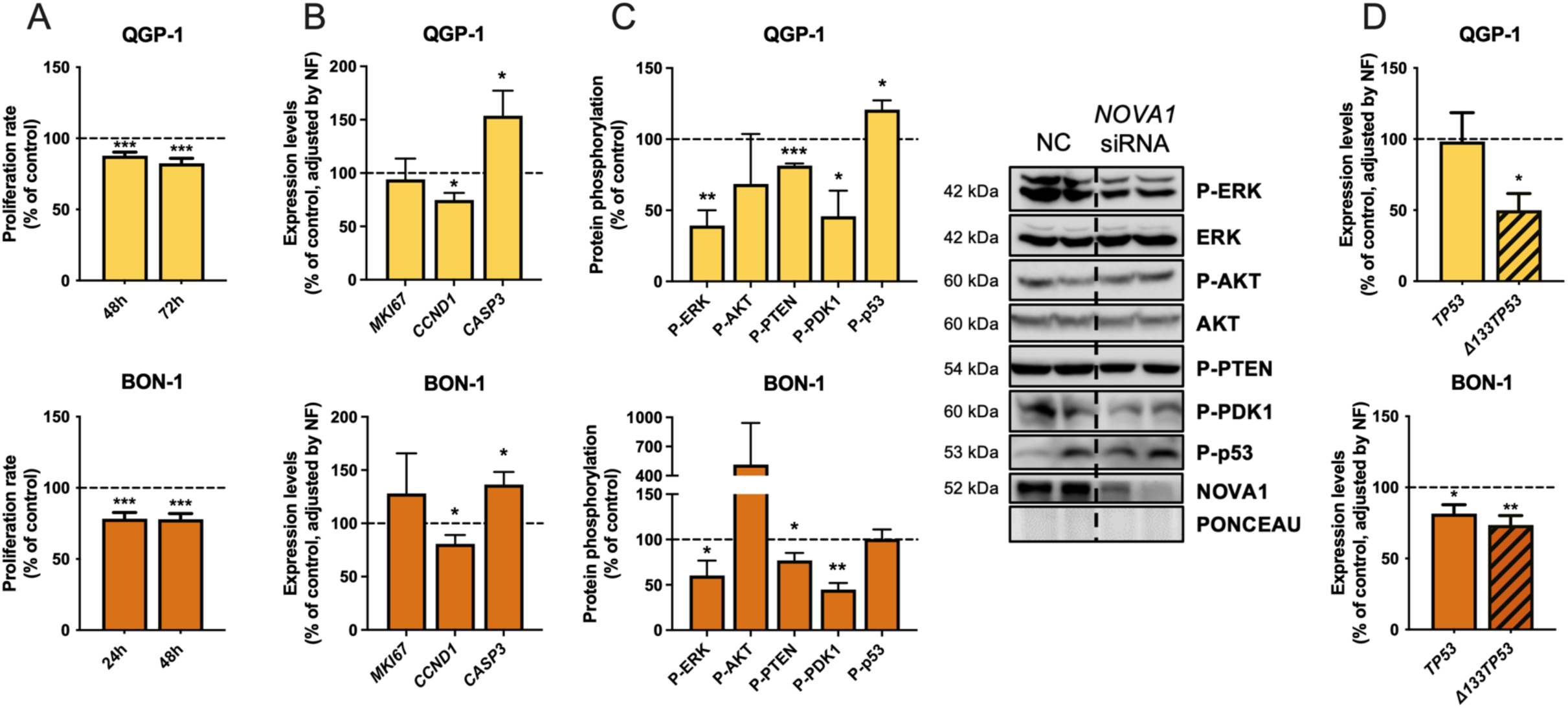
*NOVA1* silencing decreases aggressiveness features. **A**. Proliferation rate of QGP-1 (yellow; upper) and BON-1 (orange; lower) at 48 and 72 h or 24 and 48 h, respectively, after *NOVA1* silencing, compared with scramble siRNA used as control. **B**. Expression levels of mRNA of *MKI67, CCND1* and *CASP3* in QGP-1 (yellow; upper) and BON-1 (orange; lower) cell lines under *NOVA1* silencing, compared with scramble siRNA. **C**. Protein phosphorylation of ERK, AKT, PTEN, PDK-1 and p53 in QGP-1 (yellow; upper) and BON-1 (orange; lower) cell lines after *NOVA1* silencing, compared with scramble siRNA. This activation was measured by western blot and normalized with total protein or with Ponceau. **D**. *TP53* (open bars) and *Δ133TP53* (striped bars) mRNA expression levels in QGP-1 (yellow; upper) and BON-1 (orange;lower) under *NOVA1* downregulation, compared with scramble. Absolute mRNA levels were determined by qPCR and adjusted by normalization factor with *ACTB, GAPDH* and *HPRT1* housekeeping genes. Control levels were marked as a tick line. Asterisks (*, p < 0.05; **, p < 0.001; ***, p < 0.001) indicate values that significantly differ from control. In all cases, data represent mean ± SEM of n ≥ 3 independent experiments.

Then, we assessed the mRNA levels of markers commonly related to key cell functions in cancer. *NOVA1* silencing decreased, in both cell lines, the expression of *CCND1* and increased the expression of *CASP3*. In contrast, no changes were found in *MKI67*, the gene encoding the protein measured for Ki-67 index (**Fig. 4B**). Additionally, Western blotting revealed that *NOVA1* silencing decreased ERK activation, which suggests that MAPK pathways may mediate *NOVA1* actions on cell proliferation. On the other hand, *NOVA1* silencing decreased phosphorylation of both PTEN and PDK1 proteins, two pivotal mediators that play opposite roles in the activation of PI3K/AKT pathway, while AKT activation was not altered. Finally, *NOVA1* silencing increased the phosphorylation of p53 in QGP-1 but did not alter it in BON-1 (**Fig. 4C**).

The pivotal role of p53 in cancer and its distinct response to *NOVA1* silencing in the two cell types prompted us to explore this molecule in more detail. Thus, we evaluated the expression of two different isoforms of *TP53*, the gene encoding p53 protein, in order to study if *NOVA1* silencing exerts any effect on *TP53* transcription. Interestingly, we observed that this silencing clearly decreased the expression of *Δ133TP53* isoform without altering that of the canonical *TP53* in QGP-1, whereas, in contrast, both variants were significantly decreased in BON-1 (**Fig. 4D**). The silencing validation is shown in **Supplemental Figure 3B**.

### Chromatin remodeling pathway is altered under *NOVA1* downregulation

We studied how alterations in *NOVA1* may cause changes in biomarkers related with chromatin remodeling pathway. In an initial approach, we found that both *ATRX* and *DAXX* were overexpressed in our cohort of tumors, compared to non-tumoral adjacent tissue (**Fig. 5A**). In line with this, silencing of *NOVA1* led to a decrease in ATRX and DAXX protein levels in both cell lines (**Fig. 5B**). Moreover, in support of the importance of this relationship, we observed that *NOVA1* silencing decreased the expression of the oncogenic splicing variant of *TERT* gene (named as tv1) (35), without altering the expression of the canonical one (**Fig. 5C**).

**Figure 5:**
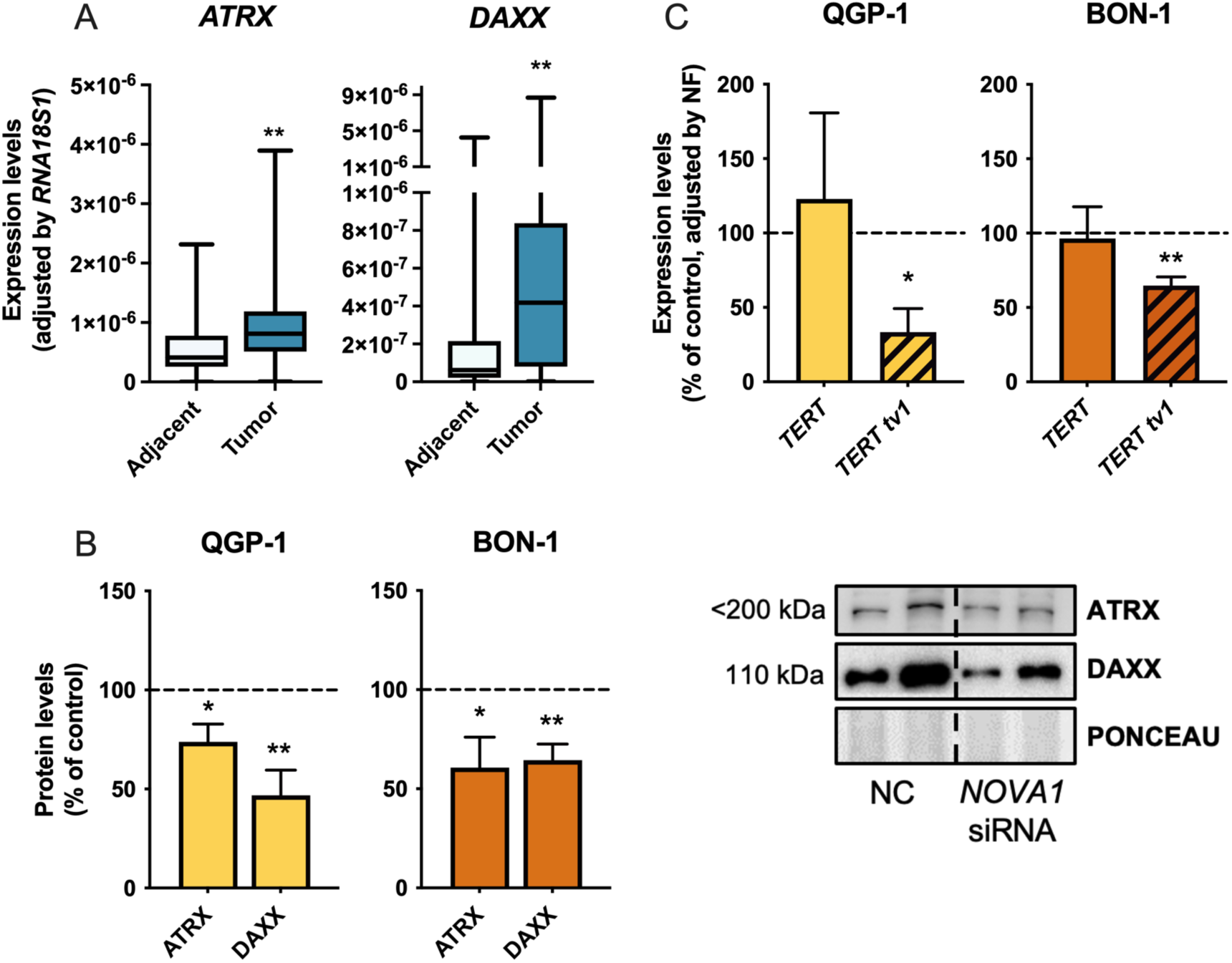
*NOVA1* silencing changes the expression and splicing of chromatin regulation genes. **A**. *ATRX* and *DAXX* mRNA expression levels in our cohort of PanNETs samples compared to non-tumoral adjacent tissue, used as control. Absolute mRNA levels were determined by qPCR and adjusted by *RNA18S1* housekeeping RNA. **B**. ATRX and DAXX protein levels after *NOVA1* silencing. Protein levels were measured by western blot, normalized with Ponceau and represented as percentage of control, marked as a tick line. **C**. Expression levels of *TERT* (open bars) and *TERT tv1* (striped bars) under *NOVA1* silencing, compared to scramble control, in QGP-1 (yellow; left) and BON-1 (orange; right). Absolute mRNA levels were determined by qPCR and adjusted by normalization factor with *ACTB, GAPDH* and *HPRT1* housekeeping genes. Asterisks (*, p < 0.05; **, p < 0.01) indicate values that significantly differ from control. In all cases, data represent mean ± SEM or median plus interquartile range of n ≥ 3 independent samples or experiments.

### Expression of *NOVA1* can influence treatment effectiveness in PanNETs

To test if *NOVA1* could influence the responsiveness of PanNETs to the main medical treatments available for this disease, *NOVA1*-silenced cells were treated with everolimus, lanreotide, octreotide and sunitinib, four currently used clinical treatments of PanNETs, as compared with scramble-silenced cells. This revealed that *NOVA1* downregulation significantly improved the antiproliferative effect of everolimus in QGP-1 cells at 72 h of treatment (**Fig. 6A**), whereas no additive effect was observed in BON-1 cell line (**Fig. 6B**). In contrast, the other treatments tested in this experimental approach did not change their effects after *NOVA1* silencing in these cells (**Supplemental Fig. 4**). These results suggest that altered *NOVA1* expression may influence selectively the response of some PanNETs to a key current treatment, everolimus.

**Figure 6.**
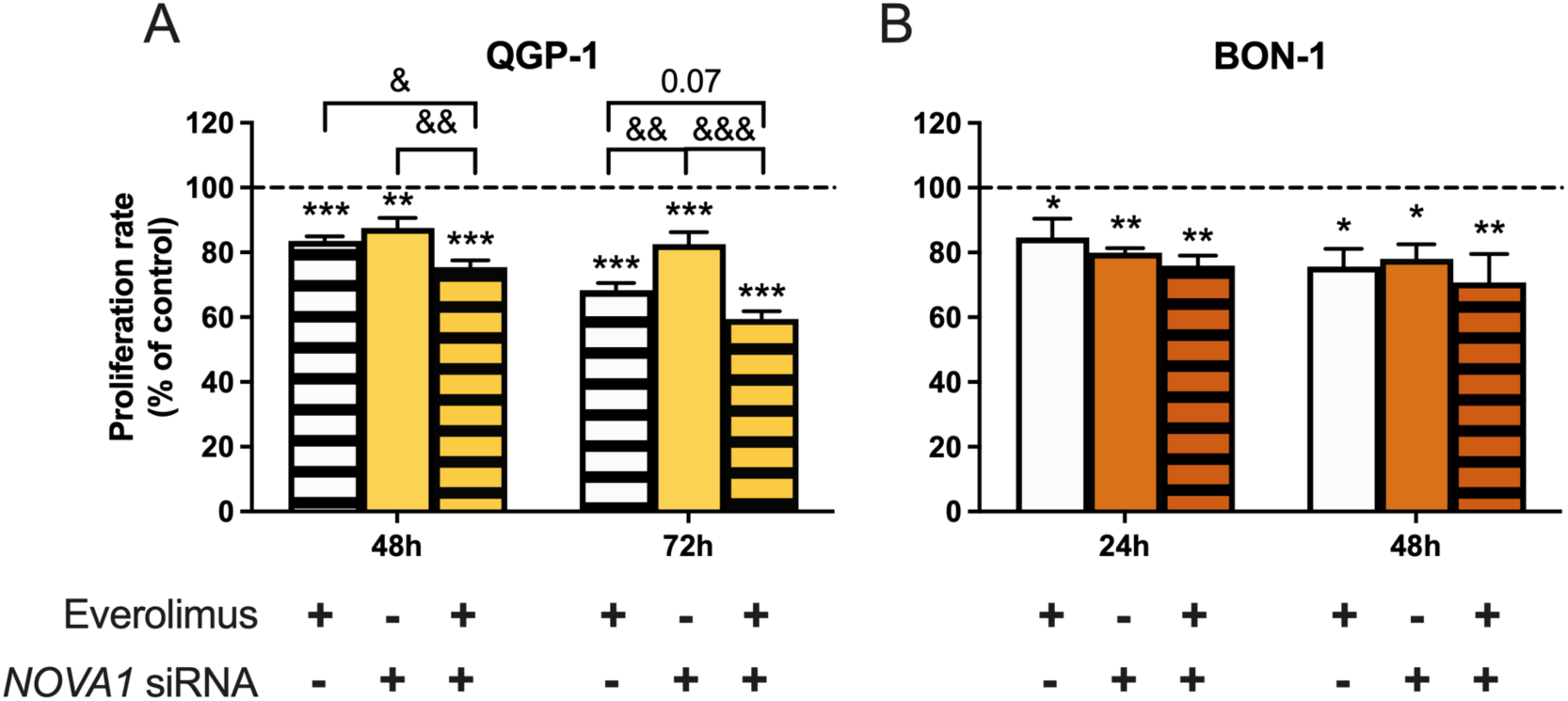
*NOVA1* silencing enhances everolimus antiproliferative effect in PanNETs cells. Proliferation rate of QGP-1 (yellow; left) and BON-1 (orange, right) cell lines after *NOVA1* silencing (color bars) plus treatment with everolimus (striped bars), compared with the non-treated scramble siRNA, used as control and marked as a tick line. Asterisks and & symbols (*/&, p < 0.05; **/&&, p < 0.01; ***/&&&, p < 0.001) indicate significant differences against the control or between groups, respectively. Data are presented as percentage of control and represent mean ± SEM of n ≥ 3 independent experiments.

## Discussion

Alteration of alternative splicing is increasingly regarded as a novel, transversal cancer hallmark, as it has been associated to many types of tumors (10, 18, 36-39). Our original discoveries of novel isoforms of SST_5_ (13, 14) and ghrelin (15) and their contribution to tumor aggressiveness in NETs (11, 12) led us to hypothesize that the splicing machinery could be involved in these events. However, a focused description of the splicing machinery had not been reported hitherto in PanNETs. Hence, our team decided to systematically characterize the pattern of expression of a representative set of components of the splicing machinery, including the core of the spliceosome and a group of selected splicing factors, as it has been recently published for pituitary NETs (23), prostate cancer (21) and glioblastoma (20).

First, comparison of the expression profile of the splicing machinery between tumor tissue and non-tumor adjacent tissue, revealed that mRNA levels of nearly 50 % of the measured genes were upregulated, whereas only one component, *ESRP2*, was found downregulated in tumor samples. Then, bioinformatic and statistical analysis of our results identified five genes, *NOVA1, PRPF8, RAVER1, SRSF5* and *SNW1*, that stand out over the rest, both because of their overexpression in virtually every single paired sample, and for their clustering ability to separate tumoral from non-tumoral samples, which was supported by ROC curves. Moreover, the possible relevance of these results was further supported by the observation that their increased levels were associated with important clinical parameters, such as vascular invasion, disease relapse or Ki-67 index, a widely used score for tumor cell proliferation with prognostic and clinical value. Among these five factors, *NOVA1* was selected for further studies based, initially, on its best fitted ROC curve (AUC > 0.86), but subsequently on a specific immunohistochemical analysis. This approach served not only to confirm at the protein level the mRNA overexpression observed in the tumor tissue, but also illustrated the intense confinement of NOVA1 immunostaining on the neuroendocrine tumor cells, as compared to the low levels present in normal endocrine cells of the non-tumoral surrounding tissue. Moreover, the close association of *NOVA1* expression levels with Ki-67 index and tumor necrosis strongly suggested that this splicing factor could be functionally linked to key PanNET cell features such as cell proliferation and death (1, 2, 40, 41). These findings were not totally unexpected in that, although *NOVA1* has classically been considered a brain-specific splicing factor, particularly relevant for neurodevelopment (42), it was also proposed earlier as a master regulator of alternative splicing in pancreatic beta cells (43), and in recent years it has also been associated to several non-neurological cancers (44, 45).

Initial *in vitro* studies using two PanNET model cell lines, BON-1 and QGP-1 cells, revealed that *NOVA1* overexpression similarly increased basal proliferation rates in both cell types. Furthermore, a preclinical model based on an immunodeficient mice with xenografted tumors indicated that BON-1 cells overexpressing *NOVA1* also displayed a higher proliferation rate than mock-transfected cells *in vivo*, thereby producing larger tumors. Correspondingly, *NOVA1* silencing decreased proliferation in both cell lines. Overall, these findings compare favorably with those recently reported in other tumors, like non-small cell lung cancer, melanoma and osteosarcoma (35, 46, 47), while the mechanisms underlying *NOVA1* actions may be tumor-specific. Thus, in PanNETs, our results reveal that *NOVA1* silencing increases ERK phosphorylation and *CCND1* mRNA levels, suggesting that it increases cell proliferation through activation of the MAPK pathway, and the subsequent involvement of cell cycle regulator *CCND1*, which are known to interact in pediatric brain tumors (48). However, the increase in *CASP3* expression suggests that an involvement of cell apoptosis should not be discarded in this context. Of note, we also discovered that *NOVA1* silencing could alter the splicing of the telomerase gene, *TERT*, by decreasing *TERT* transcript variant 1 (tv1) without altering the total expression of the gene. This alternative variant is known to exert a constitutive action that increases the length of telomeres and enhances tumor cell aggressiveness features in non-small cell lung cancer (35), in a mechanism mediated by polypyrimidine-tract binding protein 1 (*PTBP1*) that is dependent on *NOVA1* (49). Intriguingly, and possibly in relation to the above, *NOVA1* silencing also decreased protein levels of ATRX and DAXX, two genes related to chromatin remodeling and ALT, that are considered tumor suppressors, as their mutations/loss are linked to PanNET higher aggressiveness and poorer prognosis (1, 2, 7-9). Unfortunately, available knowledge on the meaning and regulation of *ATRX* and *DAXX* expression levels in PanNETs is not as advanced as that on their mutations, and the role of *TERT* similarly awaits further elucidation, beyond its limited mutational rate in well differentiated PanNETs (49). Nonetheless, our present findings may disclose an unprecedented, *NOVA1*-involving potential crosstalk between two distinct telomere length regulatory mechanisms, *TERT* and ALT, which deserves further investigation.

To gain deeper insight into the signaling pathways mediating *NOVA1* function, we next focused on those known to be linked to cell proliferation and PanNETs oncogenesis. This revealed that activation of PTEN and PDK1, two key components of the PI3K/AKT pathway, essential in PanNETs, was inhibited under *NOVA1* silencing. This finding is apparently contradictory in that these proteins are functional antagonist in the activation of this pathway, where *PTEN* is a key inhibitor and *PDK1* an important activator, closely related to *AKT* (50, 51). In fact, AKT activation itself was not altered after *NOVA1* silencing (unlike in melanoma (47)), suggesting that, to exert its actions, NOVA1 may distinctly regulate, in a tumor-specific manner, precise components of this complex signaling network, which may even play opposite roles. Interestingly, PTEN inhibition can increase cell senescence, without changes in AKT, through direct interaction with the mTOR-p53 pathway (52). Indeed, we found that *NOVA1* silencing promoted p53 activation in QGP-1 but not in BON-1, a difference that may be related to the *CDKN2A* inactivating mutation found in BON-1 (53), which would impede p53 activation, as it is its main driver in the context of senescence. In line with the above and supporting the idea that *NOVA1* silencing can activate senescence, we observed that in QGP-1 cells this silencing also downregulated selectively the *Δ133TP53* isoform, without altering full *TP53* gene expression, a relevant finding because truncated *Δ133TP53* acts as a direct inhibitor of full-length, canonical p53, especially in senescence context (54). As expected, BON-1 cells responded differently in this context, where *NOVA1* silencing inhibited the expression of both *Δ133TP53* and *TP53*. Taken together, these results suggest that the favorable actions of *NOVA1* silencing in NET cells could be exerted by increasing cell senescence, through the activation of PTEN/p53 pathway and the accompanying biasing of *TP53* transcription against the truncated *Δ133TP53* isoform.

The modulation of *NOVA1* may also entail clinically relevant implications, in that *NOVA1* silencing increased the antiproliferative effect of everolimus, an mTOR inhibitor a widely used in NETs, in QGP-1 cells, whereas no such additive effect was found in BON-1 cells. These differential, cell line-dependent results might be in consonance with our previous findings and support a role of *NOVA1* on senescence, given that, in parallel and for the same reason exposed above, the additive effect was observed in QGP-1 cells but not in BON-1 cells. Although, our present findings also suggest that the action of NOVA1 in MAPK pathway may not be independent of mTOR, because its silencing in BON-1 cells has an effect in the activation of ERK, despite not being additive to everolimus action. These complex differences between cell lines could be attributable to distinct mutations in specific components of the AKT/mTOR pathway that are differentially present in one cell line and not in the other one. This is, for example, the case of *TSC2*, one of the most important inhibitors of mTOR (55), that is mutated in BON-1 but not in QGP-1 (53), a divergence that could distinctly influence the effect of everolimus and its combined action with *NOVA1* silencing. Thus, because the set of specific mutations substantially differs in each PanNET patient and may even evolve over time in a given tumor (2, 6, 7), our results suggest that further research on *NOVA1* may guide towards the identification of novel relevant targets with therapeutic potential in a personalized manner.

When viewed together, our results reveal, for the first time, that the splicing machinery is profoundly altered in PanNETs, most of its components being upregulated. Expression of some components is associated with clinical parameters and can efficiently discriminate between tumoral and non-tumoral samples. Importantly, *in vitro* and *in vivo* studies revealed that the factor *NOVA1* can modulate proliferation and senescence in PanNETs cell lines, where it alters key signaling pathways and splicing mechanisms, and may alter the response to everolimus. These data support the splicing factor *NOVA1* as a promising candidate to develop novel biomarkers and therapeutic targets in PanNETs.

## Declarations

## Acknowledgements

We are gratefully indebted to all the patients and their families for generously donating the samples and clinical data for research purposes. We also acknowledge the technical help of the personnel of the research animal facilities and the core services of the IMIBIC (UCAIB) and University of Cordoba (SCAI).

## Availability of data and materials

The datasets used and/or analyzed during the current study are available from the corresponding author on reasonable request.

## Authors contribution

S.P.A., R.M.L, J.P.C. conceived and designed the project. S.P.A., E.A.P., R.B.E., J.M.J.V, A.C.F.F., O.R. performed the experiments. S.P.A, E.A.P., A.D.H.M., J.M.J.V., A.C.F.F., O.R., S.V., M.D.G., A.I.C., R.M.L., J.P.C. analyzed data and interpreted the results. S.P.A, R.M.L., J.P.C. prepared figures. A.D.H.M., R.S.S., R.O.S., R.S.B., M.A.G.M. acquired clinical/pathological data and samples. S.P.A., M.D.G., A.I.C., R.M.L, J.P.C. wrote the manuscript. S.P.A, E.A.P., R.B.E., A.D.H.M., J.M.J.V., A.C.F.F., O.R., S.V., R.S.S., R.O.S., R.S.B., M.A.G.M., M.D.G., A.I.C., R.M.L., J.P.C. critically revised the manuscript and approved final version.

## Ethics approval and consent to participate

All techniques carried out in this study were conducted in accordance with the ethical standards of the Helsinki Declaration, of the World Medical Association and with the approval of the University of Cordoba/IMIBIC and Ethics Committees from all the Hospitals involved in the study. Informed consent from each patient was obtained through the Andalusian Biobank (Servicio Andaluz de Salud, application code S1900499). All experimental procedures with mice were carried out following the European Regulations for Animal Care, in accordance with guidelines and regulations, and under the approval of the University/Regional’s Government Research Ethics Committees.

## List of abbreviations

NETs: neuroendocrine tumors

PanNETs: Pancreatic neuroendocrine tumors

qPCR: quantitative polymerase chain reaction

CgA: chromogranin A

mTOR: mammalian target of rapamycin

snRNAs: small nuclear RNA

FBS: fetal bovine serum

FFPE: formalin-fixed paraffin-embedded

IHC: immunohistochemistry

siRNA: small interference RNA

ROC: receiver operating characteristic

MAPK: Mitogen-activated protein kinase

PI3K: phosphatidylinositol 3-kinase.

