## Supplementary material for "Spliceosomic dysregulation unveils *NOVA1* as an actionable therapeutic target in pancreatic neuroendocrine tumors"

**A**

List of spliceosome components and splicing factors included in qPCR array.

| Major spliceosome | Minor Spliceosome | Splicing factors | | Housekeeping genes |
|---|---|---|---|---|
| *SNRNP70* | *RNU11* | *MAGOH* | *SRSF2* | *ACTB* |
| *SNRNP200* | *RNU12* | *CELF1* | *SNW1* | *HPRT* |
| *RNU4* | *RNU4ATAC* | *ESRP1* | *SND1* | *GAPDH* |
| *RNU2* | *RNU6ATAC* | *ESRP2* | *SRRM1* | |
| *RNU6* | | *NOVA1* | *SRRM4* | |
| *U2AF1* | | *SFPQ (PSF)* | *SRSF3* | |
| *U2AF2* | | *PTBP1 (PTB)* | *SRSF4* | |
| *SF3B1* | | *RAVER1* | *SRSF5* | |
| *SF3B1 tv1* | | *RBM17* | *SRSF6* | |
| *TCERG1* | | *RBM3* | *SRSF9* | |
| *PRPF40A* | | *RBM45* | *TIA1* | |
| *PRPF8* | | *KHDRBS1 (SAM68)* | *TRA2A* | |
| *RBM22* | | *SRSF1* | *TRA2B* | |
| | | *SRSF10* | | |

**B**

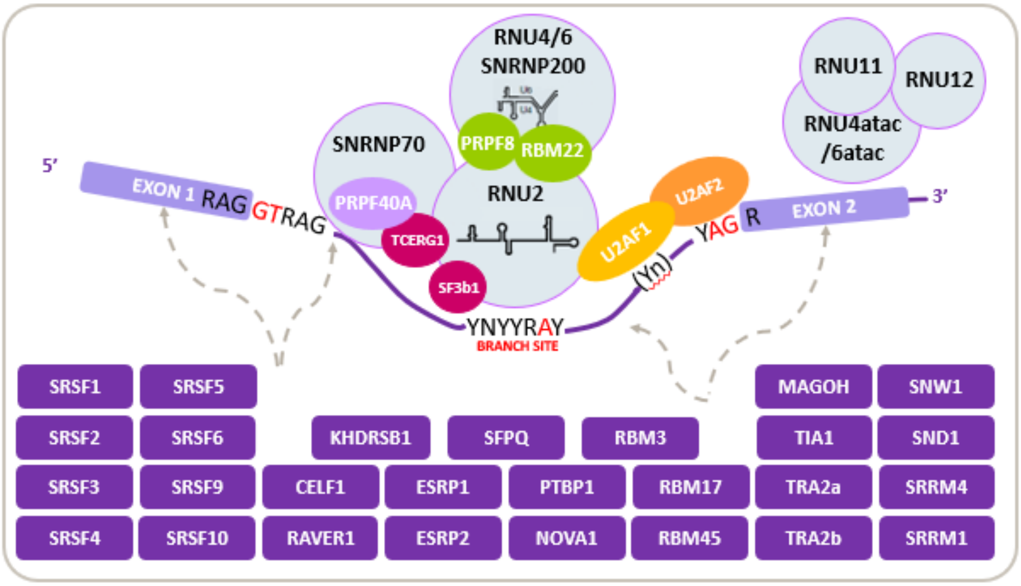


**Supplemental Figure 1.** List (**A**) and schematic representation (**B**) of the splicing machinery components included in the qPCR array analysis of this study.

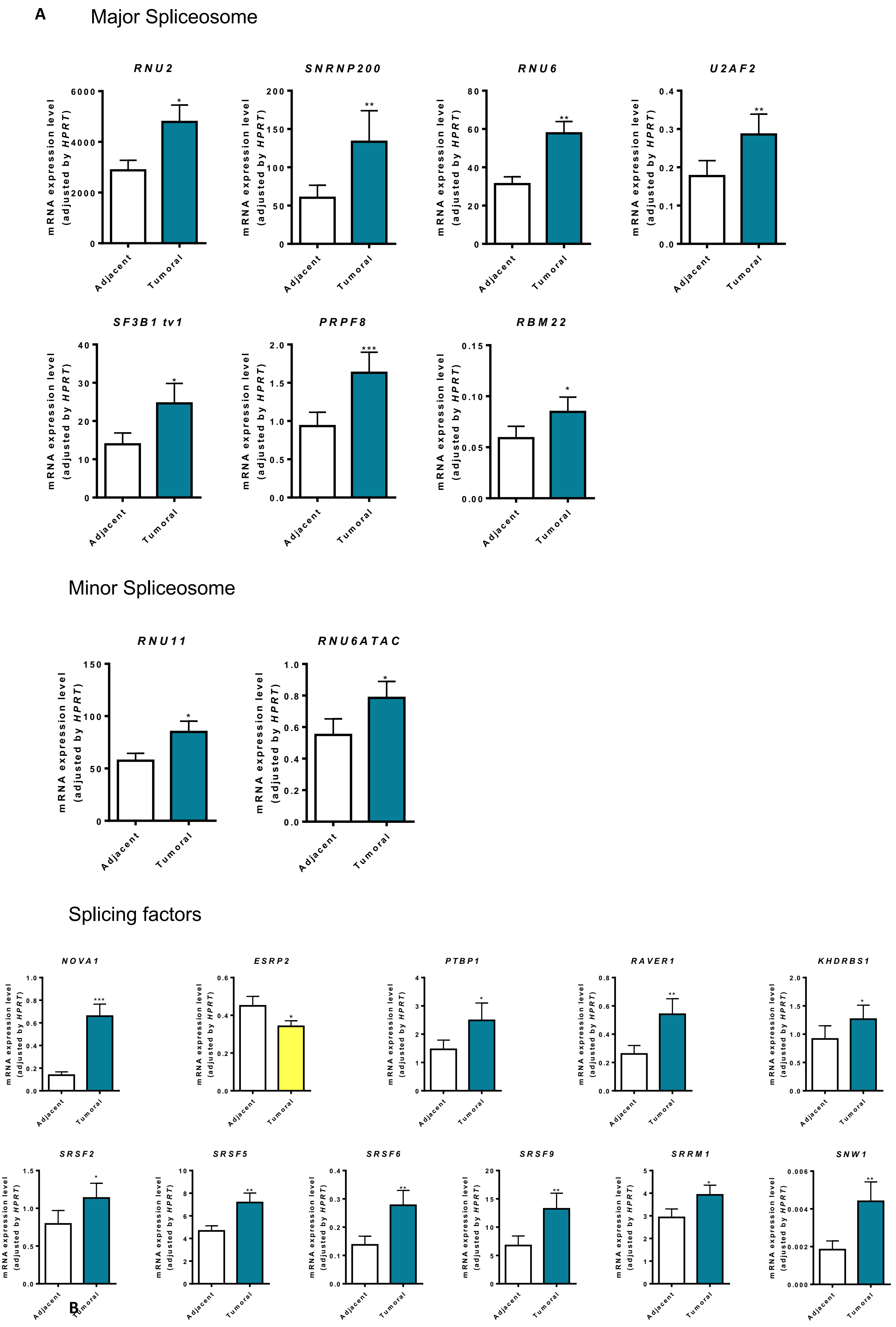
A
Major Spliceosome
RNU2
SNRNP200
RNU6
U2AF2
SF3B1 tv1
PRPF8
RBM22
Minor Spliceosome
RNU11
RNU6ATAC
Splicing factors
NOVA1
ESRP2
PTBP1
RAVER1
KHDRBS1
SRSF2
SRSF5
SRSF6
SRSF9
SRRM1
SNW1
mRNA expression level (adjusted by HPRT)
Adjacent
Tumoral
B

### Major Spliceosome

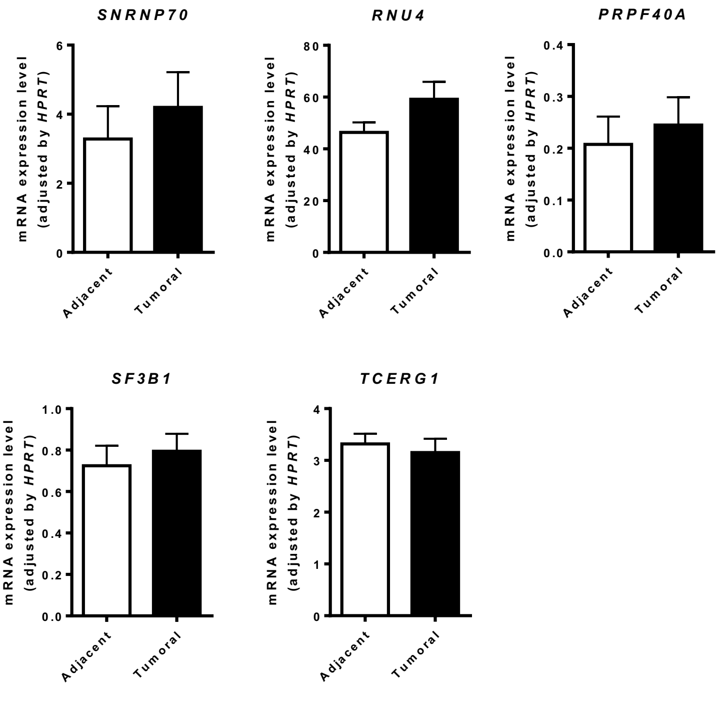


### Minor Spliceosome

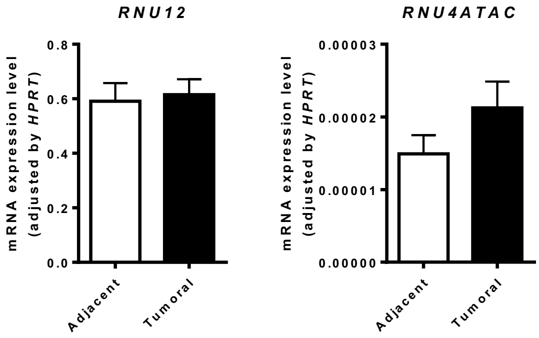


### Splicing factors

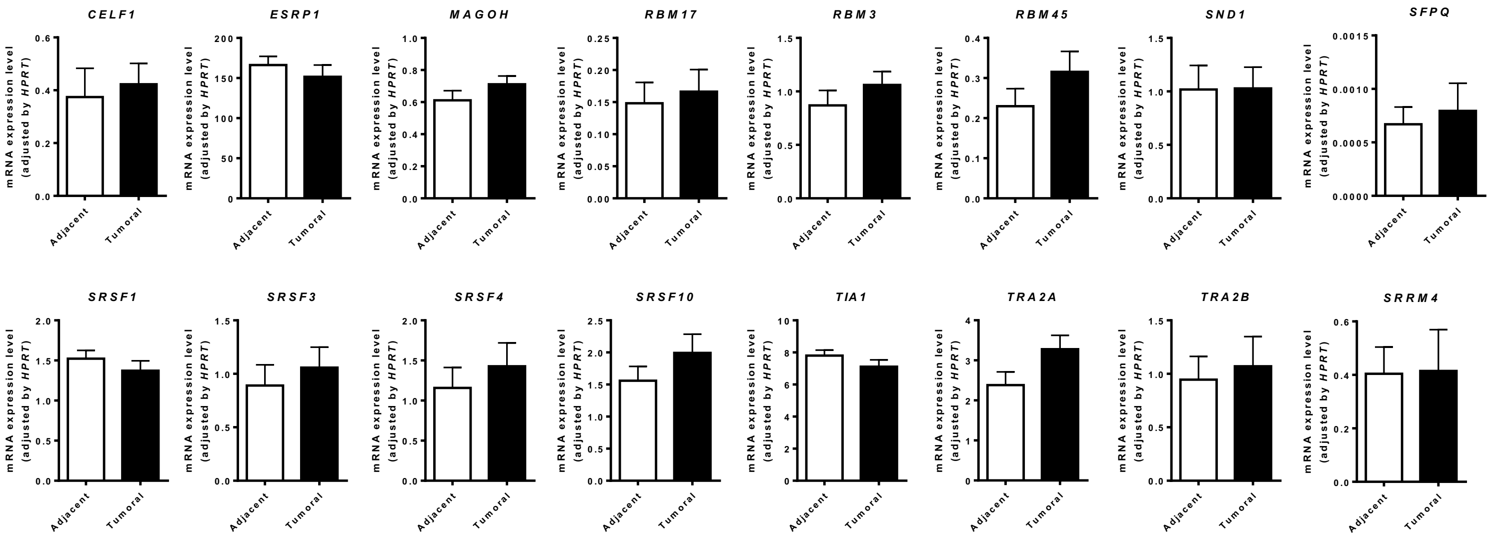

**Supplemental figure 2.** Differential mRNA expression levels of splicing machinery components in PanNETs tumoral samples compared with non-tumoral adjacent tissue. The components with significant changes (**A**) are in the upper panel and those without significant alterations (**B**) are in the bottom panel.

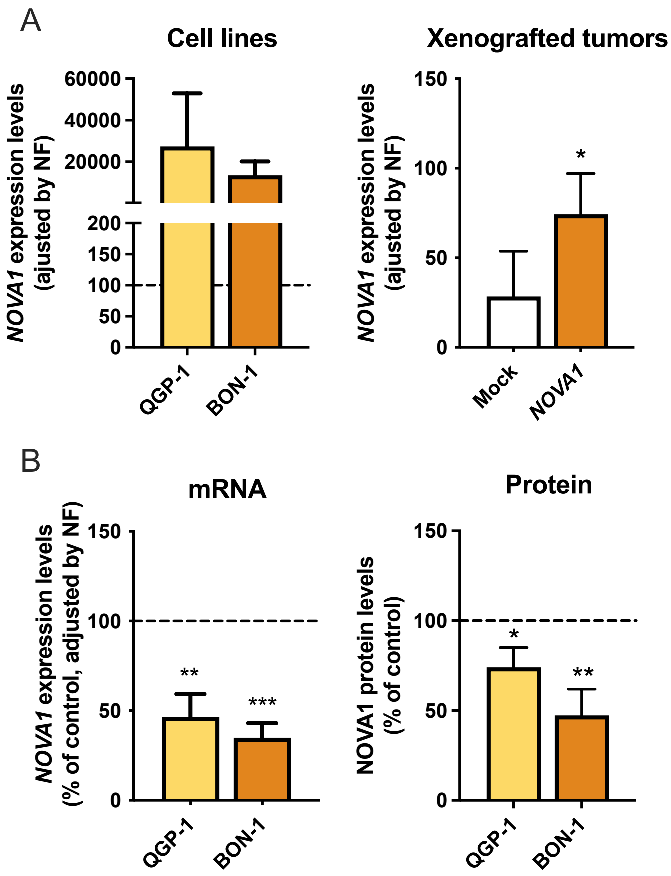


**Supplemental figure 3. A.** *NOVA1* overexpression validation by qPCR in cell lines and xenografted tumors after euthanasia. **B**. *NOVA1* silencing validation in mRNA and protein expression levels.

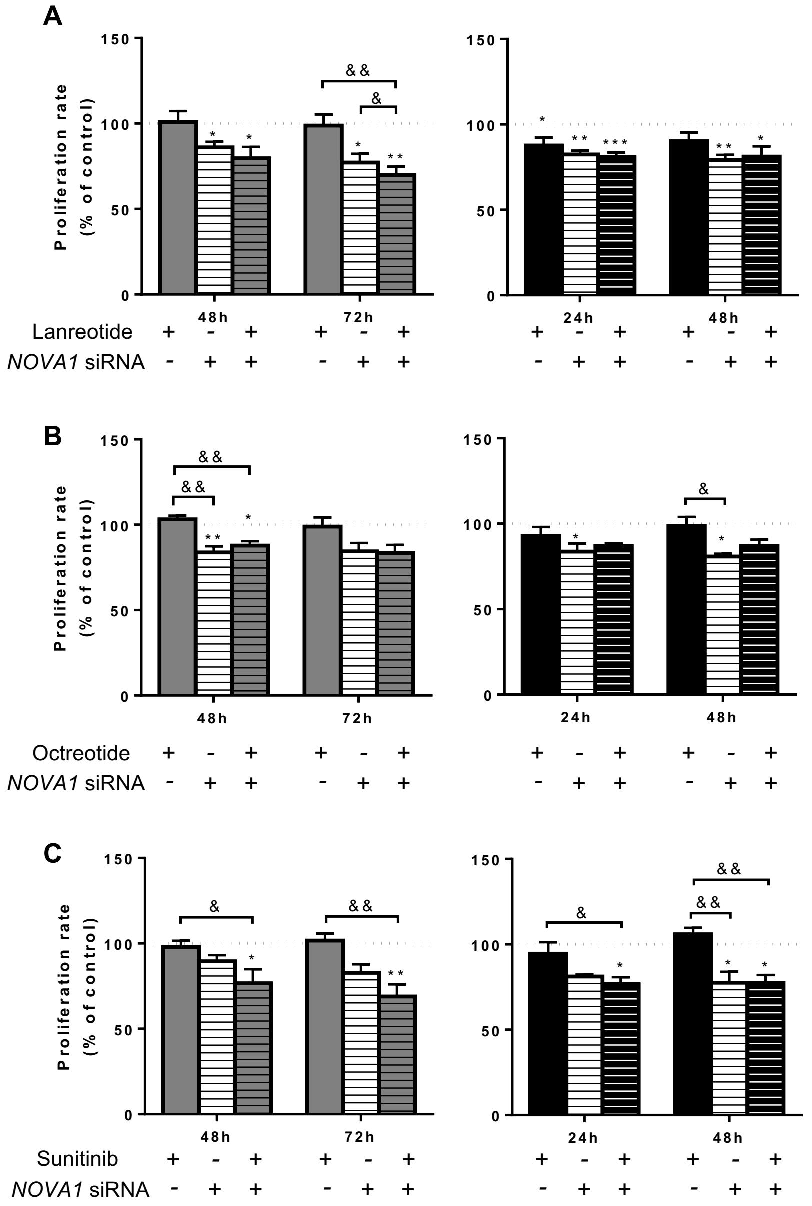


**Supplemental figure 4.** Proliferation rate with combination of *NOVA1* silencing with PanNETs classical treatments: lanreotide (**A**), octreotide (**B**) and sunitinib (**C**), in QGP-1 (grey, left) and BON-1 (black, right). Asterisks and & symbols (*, $p < 0.05$; **, $p < 0.01$; ***, $p < 0.001$) indicate significant differences. In all cases, data represent mean ± SEM of $n \geq 3$ independent experiments.

| | Gene | Primer Sequence (Sense, Se) | Primer Sequence (Antisense, As) | Product Size | Access Number |
|---|---|---|---|---|---|
| Splicing Factors | *CELF1* | AACAGAAGAGAATGGCCCAGC | TGCTGAAGGAGTGCTAAATACTG | 121 | NM_006560.3 |
| | | | | | NM_198700.2 |
| | *ESRP1* | TTTTGGGATCACTGCTGGGG | TGTCCCACCTTCTTGTTGGC | 108 | NM_017697.3 |
| | | | | | NM_001034915.2 |
| | | | | | NM_001122826.1 |
| | | | | | NM_001122825.1 |
| | | | | | NM_001122827.1 |
| | *ESRP2* | AGAGCCCAGCAGTCAATTGTT | GTCTCACTGTCCACCACATCAG | 96 | NM_024939.2 |
| | *KHDRBS1* | GAGCGAGTGCTGATACCTGTC | CACCAGTCTCTTCCTGCAGTC | 106 | NM_006559.2 |
| | | | | | NR_073498.1 |
| | | | | | NR_073499.1 |
| | *MAGOH* | GCCAACAACAGCAATTACAAGA | TTATTCTCTTCAGTTCCTCCATCAC | 88 | NM_002370.3 |
| | *NOVA1* | TACCCAGGTACTACTGAGCGAG | CTGGTTCTGTCTTGGCCACAT | 124 | NM_002515.2 |
| | | | | | NM_006489.2 |
| | | | | | NM_006491.2 |
| | *PTBP1* | TGGGTCGGTTCCTGCTATT | CAGATCCCCGCTTTGTAC | 111 | NM_002819.4 |
| | | | | | NM_031990.3 |
| | | | | | NM_031991.3 |
| | *RAVER1* | GTAACCGCCGCAAGATACTG | CGAAGGCTGTCCCTTTGTATT | 126 | NM_133452.2 |
| | *RBM17* | CAAAGAGCCAAAGGACGAAA | TACATGCGGTGGAGTGTCC | 107 | NM_032905.4 |
| | | | | | NM_001145547.1 |
| | *RBM3* | AAGCTCTTCGTGGGAGGG | TTGACAACGACCACCTCAGA | 98 | NM_006743.4 |
| | *RBM45* | CCCATCAAGGTTTTCATTGC | TTCCCGCAGATCTTCTTCTG | 123 | NM_152945.3 |
| | *SFPQ* | TGGTAGGGGGTGAAAGTG | TTAAAAACAAGAAATGGGGAAATG | 125 | NM_005066.2 |
| | *SND1* | ACTACGGCAACAGAGAGGTCC | GAAGGCATACTCCGTGGCT | 101 | NM_014390.3 |
| | *SNW1* | ATGCGTGCCCAAGTAGAGAG | TCCCCATCCTCTTTTTCCA | 134 | NM_001318844.1 |
| | | | | | NM_012245.2 |
| | *SRRM1* | GTAGCCCAAGAAGACGCAAA | TGGTTCTGTGACGGGGAG | 108 | NM_001303448.1 |
| | | | | | NM_005839.3 |
| | | | | | NM_001303449.1 |
| | *SRRM4* | CCTTCACCACCTCCTCAC | TTCGGCACATTCCAGACA | 113 | NM_194286.3 |
| | *SRSF1* | TGTCTCTGGACTGCCTCCA | TGCCATCTCGGTAAACATCA | 98 | NM_006924.4 |
| | | | | | NM_001078166.1 |
| | *SRSF10* | CTACACTCGCCGTCCAAGAG | CCGTCCACAAATCCACTTTC | 103 | NM_006625.5 |
| | | | | | NM_054016.3 |
| | | | | | NM_001191005.2 |
| | | | | | NM_001191006.2 |
| | | | | | NM_001191007.2 |
| | | | | | NM_001191009.2 |
| | | | | | NM_001300936.1 |
| | | | | | NM_001300937.1 |
| | *SRSF2* | TGTCCAAGAGGGAATCCAAA | GTTTACACTGCTTGCCGATACA | 113 | NM_003016.4 |
| | | | | | NM_001195427.1 |
| | *SRSF3* | TAACCCTAGATCTCGAAATGCATC | CATAGTAGCCAAAAGCCCGTT | 117 | NM_003017.4 |
| | | | | | NR_036610.1 |
| | *SRSF4* | GGAACTGAAGTCAATGGGAGAA | CTTCGAGAGCGAGACCTTGA | 110 | NM_005626.4 |
| | *SRSF5* | GCAAAAGGCACAGTAGGTCAA | TTTGCGACTACGGGAACG | 92 | NM_001039465.1 |
| | | | | | NM_006925.4 |
| | | | | | NM_001320214.1 |
| | *SRSF6* | AGACCTCAAAAATGGGTACGG | CTTGCCGTTCAGCTCGTAA | 82 | NM_006275.5 |
| | | | | | NR_034009.1 |
| | *SRSF9* | CCCTGCGTAAACTGGATGAC | AGCTGGTGCTTCTCTCAGGA | 87 | NM_003769.2 |
| | *TIA1* | TAAATCCCGTGCAACAGCAGA | TATGCAGGAACTTGCCAACCA | 124 | NM_022037.2 |
| | | | | | NM_022173.2 |
| | *TRA2A* | TCAAAGGAGGCTATGGAAAGG | TGTGTGCGCTCTCTTGGTTA | 90 | NM_013293.4 |
| | | | | | NM_001282757.1 |
| | | | | | NM_001282758.1. |
| | | | | | NM_001282759.1 |
| | *TRA2B* | GATGATGCCAAGGAAGCTAAAG | AGGTAGGTCTCCCCATGTAAATTC | 130 | NM_004593.2. |
| | | | | | NM_001243879.1 |

| | | | | | |
|---|---|---|---|---|---|
| Spliceosome Components | *PRPF40A* | GCTCGGAAGATGAAACGAAA | TGTCCTCAAATGCTGGCTCT | 130 | NM_017892.3 |
| | *PRPF8* | TGCCCACTACAACCGAGAA | AGGCCCGTCCTTCAGGTA | 139 | NM_006445.3 |
| | *RBM22* | CTCTGGGTTCCAACACCTACA | GGCACAGATTTTGCATTCCT | 137 | NM_018047.2 |
| | *SNRNP70* | TCTTCGTGGCGAGAGTGAAT | GCTTTCCTGACCGCTTACTG | 114 | NM_001301069.1 |
| | *RNU11* | AAGGGCTTCTGTCGTGAGTG | CCAGCTGCCCAAATACCA | 108 | NR_004407.1 |
| | *RNU12* | ATAACGATTCGGGGTGACG | CAGGCATCCCGCAAAGTA | 106 | NR_029422.1 |
| | *RNU2* | CTCGGCCTTTTGGCTAAGAT | TATTCCATCTCCCTGCTCCA | 116 | NR_002716.3 |
| | *RNU4* | TCGTAGCCAATGAGGTCTATCC | AAAATTGCCAGTGCCGACTA | 103 | NR_003925.1 |
| | *RNU4ATAC* | GTTGCGCTACTGTCCAATGA | CAAAAATTGCACCAAAATAA | 85 | NR_023343.1 |
| | *RNU6* | CGCTTCGGCAGCACATATA | AAAATATGGAACGCTTCACGAA | 101 | NR_004394.1<br>NR_125730.1<br>NR_104084.1<br>NR_104088.1<br>NR_104080.1 |
| | *RNU6ATAC* | TGAAAGGAGAGAAGGTTAGCACTC | CGATGGTTAGATGCCACGA | 112 | NR_023344.1 |
| | *SF3B1* | CAGTTCCGTCTGTGTGTTCG | GCTGCCTTCTTGCCTTGA | 101 | NM_012433.3<br>NM_001005526.2<br>NM_001308824.1 |
| | *SF3B1 tv1* | GCAGACCGGGAAGATGAATA | TTTTCCCTCCATCTGCAAAA | 88 | NM_012433.3 |
| | *SNRNP200* | GGTGCTGTCCCTTGTTGG | CTTTCTTCGCTTGGCTCTTCT | 103 | NM_014014.4 |
| | *TCERG1* | GAGGAGCCCAAAGAAGAGGA | CACCAGTCCAAACGACACAC | 112 | NM_006706.3<br>NM_001040006.1 |
| | *U2AF1* | GAAGTATGGGGAAGTAGAGGAGATG | TTCAAGTCAATCACAGCCTTTTC | 120 | NM_006758.2<br>NM_001025203.1<br>NM_001025204.1 |
| | *U2AF2* | CTTTGACCAGAGGCGCTAAA | TACTGCATTGGGGTGATGTG | 130 | NM_007279.2<br>NM_001012478.1 |

| Housekeeping Genes | | | | | |
|---|---|---|---|---|---|
| | *ACTB* | ACTCTTCCAGCCTTCCTTCCT | CAGTGATCTCCTTCTGCATCCT | 176 | NM_001101 |
| | *GAPDH* | AATCCCATCACCATCTTCCA | AAATGAGCCCCAGCCTTC | 122 | NM_002046 |
| | *HPRT1* | CTGAGGATTTGGAAAGGGTGT | TAATCCAGCAGGTCAGCAAAG | 157 | NM_000194.2 |

| Isoforms | | | | |
|---|---|---|---|---|
| *TP53* | AAGGAAATTTGCGTGTGGAG | CCAGTGTGATGATGGTGAGG | 180 | NM_001276760.2 |
| *ATRX* | TGTTTTCAGCCAGTCCCTCA | GCCACTTCCCCTCACCTTTA | 118 | NM_000489.5 |
| *DAXX* | AAGCCTCCTTGGATTCTGGT | CTGCTGCTGCTTCTTCCTCT | 237 | NM_001141969.2 |
| *MKI67* | GACATCCGTATCCAGCTTCCT | GCCGTACAGGCTCATCAATAAC | 139 | NM_002417 |
| *CCND1* | CCTCGGTGTCCTACTTCAAAT | TCCTCCTCGCACTTCTGTTC | 108 | NM_053056.2 |
| *CASP3* | TTTTTCAGAGGGGATCGTTG | GTCTCAATGCCACAGTCCAGT | 97 | NM_001354777.1 |

**Supplemental table 1.** List of primers used in this study, including the name of the gene, the sequence of the primers, the length of the amplicon and the NCBI reference.

| Random Forest | AUC |
|---|---|
| *RNU6 SF3B1 tv1 TRA2B ESRP2 SRSF9 RAVER1 PRPF8 SND1* | 0,887 |
| *NOVA1 RNU6 SF3B1 tv1 TRA2B ESRP2 KHDRBS1 RAVER1 PRPF8* | 0,879 |
| *NOVA1 RNU6 SF3B tv1 TRA2B ESRP2 KHDRBS1 RAVER1 PRPF8 RNU11* | 0,878 |
| *SF3B1 tv1 TRA2B ESRP2 SRSF9 PRPF8* | 0,877 |
| *RNU6 SF3B tv1 TRA2B ESRP2 SRSF9 RAVER1 PRPF8* | 0,873 |

| Logistic Regression | AUC |
|---|---|
| *TRA2B ESRP2 SNW1 SRSF9 SRSF5 SF3B1 SRRM1 RNU5 RBM3* | 0,897 |
| *SF3B1 tv1 TRA2B ESRP2 SRSF9 U2AF2 CELF1 SFPQ SF3B1* | 0,897 |
| *SF3B1 tv1 TRA2B ESRP2 SRSF9 U2AF2 CELF1 SFPQ* | 0,893 |
| *SF3B1 tv1 TRA2B ESRP2 SRSF9 U2AF2 CELF1 SFPQ SF3B1 PTBP1* | 0,893 |
| *SF3B1 tv1 TRA2B ESRP2 SRSF9 U2AF2 CELF1 SFPQ TCERG1* | 0,891 |

**Supplemental table 2.** Random forest (upper panel) and simple logistic regression (lower panel) analyses with mRNA expression data of splicing machinery components in our cohort of PanNETs, which highlights the genes with better clustering features. AUC represents Area Under the Curve of these analyses.
